# Bistablity in Fluorescence from a purple non-sulfur bacteria

**DOI:** 10.1101/132498

**Authors:** Anirban Bose, Sufi O Raja, Sanhita Ray, Anjan Kr Dasgupta

## Abstract

Bistable optical emission has been observed for photosynthetic purple non-sulfur bacteria *Rhodobacter capsulatus* SB1003. The microbes respond to UV excitation (at 395nm) in a bifurcating way one branch corresponding to increase and the other corresponding to diminishing fluorescent emission in the range 590-685nm.The switching between such bifurcating branches can be observed when parameters like concentration, temperature are varied or static magnetic field is applied. Thus switching from amplification to reduction occurs if fluorophore concentration lowered. Again if temperature is lowered a steady quenching (instead of amplification) of fluorescence is observed. However presence of magnetic field of the order of 0.5 T reverts this and once again the systems resumes its fluoresence amplifying state. We propose that aggregation of bacterial porphyrin and regulation of such aggregation by photon excitation may explain this bistablity. Possible ecological implication of the photosynthetic bistability is suggested.

## Introduction

Bistability, is a concept implying a duality of steady states, a switchability of one from the other, appeared in biological literature in various forms (1). Broadly the term expresses existence of non-unique steady states, this in turn implying a possible dynamical switching from one to the other in response to a mild parametric change. Thus in genetics, epigenetics (2)and microbial ecology (3) bistability appears in order to describe existance of a dual genomic or ecological equilibria. It also appears in the bio-geosphere in sudden switching of oxygen concentration during the evolution of the bio-geosphere (4). There was obviously a sudden surge of photosynthetic activity. Bistability in photochemical processes have been reported earlier in several photochromic compounds, polymers and derivatives (5–8).

In spite of the presence of such a vast literature bistability has not been reported for a natural photosynthetic or cyno-bacteria. In general we find different classes of fluorescent biomolecules that are (9, 10) mostly susceptible to photo-bleaching. Here we show bistability in fluorescence emission from *Rhodobacter capsulatus* SB1003, the switchability depending on concentration and temperature. The work was inspired by our preliminary observation that we made while studying the fluorescence recovery after photobleaching (FRAP) experiment (11). FRAP is based on the high propensity of bleaching by fluorophores and its recovery by diffusion of unbleached molecules. Our study shows the fluorescence from *Rhodobacter capsulatus* SB1003 became brighter instead of bleaching, under a confocal microscope during relatively prolong UV excitation.

Photosynthetic purple non-sulfur bacteria *Rhodobacter capsulatus* SB1003 is one of the most primitive photoheterotroph that contain photoreceptors in the form of porphyrin derivatives like non-covalently bound cofactors, bacterio-chlorophyll and bacterio-pheophytin in the reaction centres of their photosynthetic membrane complexes (12). This class of α-Proteobacteria has been employed as a model system for studying photosynthesis (13) and different classes of porphyrin production under various growth conditions (14). Therefore bistability of the photosynthetic machinery of this purple bacteria seems to have important ecological and evolutionary implications.

These bacteria possess behavioral competencies to respond external stimuli by altered electron transport system which modulate their intra-cellular energy level to obtain optimal metabolic activity known as energy-taxis (15). Such behavioral response is very essential survival strategy for obtaining highest metabolic yields at different external energy levels. The photosynthetic microbes are of particular interest as they induce a thermodynamically favorable condition for supporting the ecological network of different life forms by contributing negative entropy in the form of photon capture (16). The general conjecture is that a sensory perception for species like *Rhodobacter* can respond to external light stimulus through electron transport system in their photosynthetic apparatus (17). Numerous models have been reported where the role of microbes in overall metabolic energy transfer circuitry of different ecological niche have been emphasized (18–21).

Notable point that we report in the present paper is that spin perturbations agents like static magnetic field (22–24) can play important role in this bistable emission. Possible roles of porphyrin aggregate formation with increasing temperature, concentration and interaction with static magnetic field effect have been investigated that could help one to truly embed the photon utilization and regulation machinery in designing and device level implementation of artificial photosynthesis.

## Material and Methods

All chemical reagents were used of analytical grade Sigma-Aldrich (USA) or SRL (India) products.

### Bacterial Growth

The bacteria *Rhodobacter capsulatus* strain SB1003 was a gift from Dr. Patrick Hallenbeck, Department of Microbiology and Immunology, University of Montreal, Canada. For experimental purposes *Rhodobacter capsulatus* SB1003 were grown photoheterotrophicaly in yeast extract supplemented RCV medium (4g Malic acid, 1g NH_4_Cl, 120 mg MgCl_2_, 75 mg CaCl_2_, 20 mg Sodium EDTA salt, 1mg Nicotinic acid, 10 mM KPO_4_ buffer, 1000 ml dH_2_O supplemented with 0.3g Yeast extract powder, _*P*_H 6.8 +0.2). The culture was incubated at room temperature under continuous illumination for 6-8 days(25) in screw cap bottles. The full grown bacterial culture and the cell free extract were used for monitoring fluorescence properties.

### Live Cell Imaging

Microscopic fluorescence study was initially done using bacterial biofilm (26) in an inverted confocal microscope (Olympus, FV 1000). A 405 nm laser was used as excitation source and fluorescence emission was observed between 570-670 nm. Time lapse imaging of the samples were acquired using continuous laser at 1 minute time interval for a period of nine minutes. Further study was carried out with full intensity laser for a period of one minute.

### Basic Spectroscopic Studies

Spectroscopic measurements of samples were obtained by scan method setting, absorption range from 250 nm to 1000 nm, at 1 nm bandwidth and 600 nm/min scan speed using Thermo Scientific *™* Evolution 300 UV-Vis spectrophotometer. The fluorescence properties of samples were studied with Photon Technology International (PTI) fluorimetric setup (Quantamaster *™*40). A 72 W Xenon lamp was used as an illumination source and the detection was preceded by passing the emitted beam through an optical chopper and emission monochromator. The slit width was set at 4 nm for both excitation and emission. For temperature dependent fluorescence measurement a software controlled Peltier was employed. The magnetic field application is illustrated in figure (S1) of supplementary. A small square shaped magnet (field strength 295mT) was placed under the quartz cuvette, in the cuvette-holder of above mentioned fluorimetric set up to study the effect of static magnetic field (SMF).The temperature variation experiment in presence and absence of SMF was performed using this setup.

### Determination of kinetic parameters: Spectroscopic Data Analysis

Graphs were plotted considering the initial fluorescence amplification rate along Y axis and other variables such as temperature or concentration along the X axis. Change in fluorescence intensity (count) with per unit time was expressed by a first order kinetic parameter k (*s*^−1^). In case of amplification, k value is positive. A negative K value could be obtained if only bleaching occurs. To obtain k, only the initial rate was evaluated using first 20 seconds of data. Solution to the first order equation 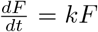, representing the rate of fluorescence intensity (F) change. Matlab (Mathworks USA) was used to obtain the best value of k. Multiple k values were determined from 5 independent experiments using log linear regression and this was followed by a box plot representation of data.

### Fluorescence Life Time Measurement

Fluorescence lifetime measurements were performed using 370nm sub-nanosecond pulsed LED source from HORIBA Scientific Single Photon Counting Controller: FluoroHub. Fluorescence Lifetime histogram was obtained using MicroTime 200, PicoQuant GmbH. Detail description of the instrument is given in the experimental set up section of Single Molecule Spectroscopy.

### Size Measurement of cell free extract by Dynamic Light Scattering (DLS)

400 *μ*L of sample was used for DLS measurement (Zetasizer Nano Series; Malvern Instruments, Malvern, UK) at 278K, 288K, 298K, 303K and 313K,respectively. Effect of a weak magnetic field on sample at 278K was studied by placing the magnet just below the cuvette during the measurement.

### Experimental Set up

A square shaped magnet of 295mT was placed under the cuvette each time to study the effect of SMF during the measurements (Figure S1).

### Organic extraction

The spectral behavior of the cell free extract was found to be changed upon solvent perturbation (Figure (S3)). When acetone-methanol (7:2) was selected as solvent medium two additional peaks appeared at 594 nm and 635 nm similar to Coproporphyrin and Mg-Protoporphyrin monoethyl ester fluorescence (27). Notably, the excitation is also red shifted to 411nm and 403 nm for the respective emissions. The split of the original spectral pattern (see figure(S2a)) into two peaks reveals the fact that the integrity of the membrane bound complex is sensitive to solvent perturbation. Notably, only one peak (635nm) among the two, is amplified with time when there is a prolonged excitation (see dotted profile in the figure(S3).

### SDSPAGE

The Fluorescence property of the photosynthetic membrane complex was revealed when the SDS-PAGE was illuminated by UV trans-illuminator (see figure (S4)). In this figure, each of the panels shows electrophoresis pattern of the different bacterial cell membrane fractions. The bands are visible even without illumination because of intrinsic color and the presence of convergent illuminated bands imply that the colored fractions are fluorescent. In the upper panel of figure(S4), lane A and lane D consist of FPLC fractions; lane B and lane C consist of membrane fractions obtained after ultracentrifugation. In the lower panel, all the lanes (B1,B2,B3,B4) represent the ultra-centrifuged fractions. The notable point is, in the left panel the color of the bands represent the membrane bound photosynthetic complexes (in absence of any conventional staining), and in the right panel the fluorescence is observed in the same position of the gel (during exposure to a UV transilluminator).

### Atomic Force Microscopy (AFM)

Glass cover slips were properly cleaned and incubated in sample solution for half an hour. Then the cover glass pieces were dried and analyzed by AFM (Veeco multi mode NanoScope IIIa) with tapping mode with a tip model RTESPA equipped with 110 O-cm phosphorous (n)-doped Si at scanning rate of 1 Hz using a phase data type and a resonant frequency of 314.5 kHz. In the figure(S5), the upper panel shows AFM micrograph image of cell free extract and the lower panel shows 3D representation of cell free extract and bacterial culture grown with excess magnesium supplement respectively (Left to right).

## Results

### Confocal microscopy

The microscopic data described in figure(1) reveals that whole bacterial cells showed enhanced fluorescence emission over a longer time period up to 10 minutes. The panels numbered 1,2,…,10, corresponds to minutes of excitation exposure of the sample. When a FRAP (Fluorescence Recovery After Photo-bleaching) experiment was carried out with bacterial cells, the fluorescence emission intensity gradually increased instead of photo-bleaching. The behavior is similar to what is found in figure(S2a) of supplementary information. For further study the intensity of the incident laser (405 nm) beam was increased from 25%to 100% and the sample was exposed for one minute. The result is shown in the lower panel of figure(1). The indices A and B refer to per-excitation and post-excitation states of the sample. This time there was a dual effect, the central part showing a bleaching (see panel B of figure(1) and the peripheral region showing an intense fluorescence.

**Figure 1:**
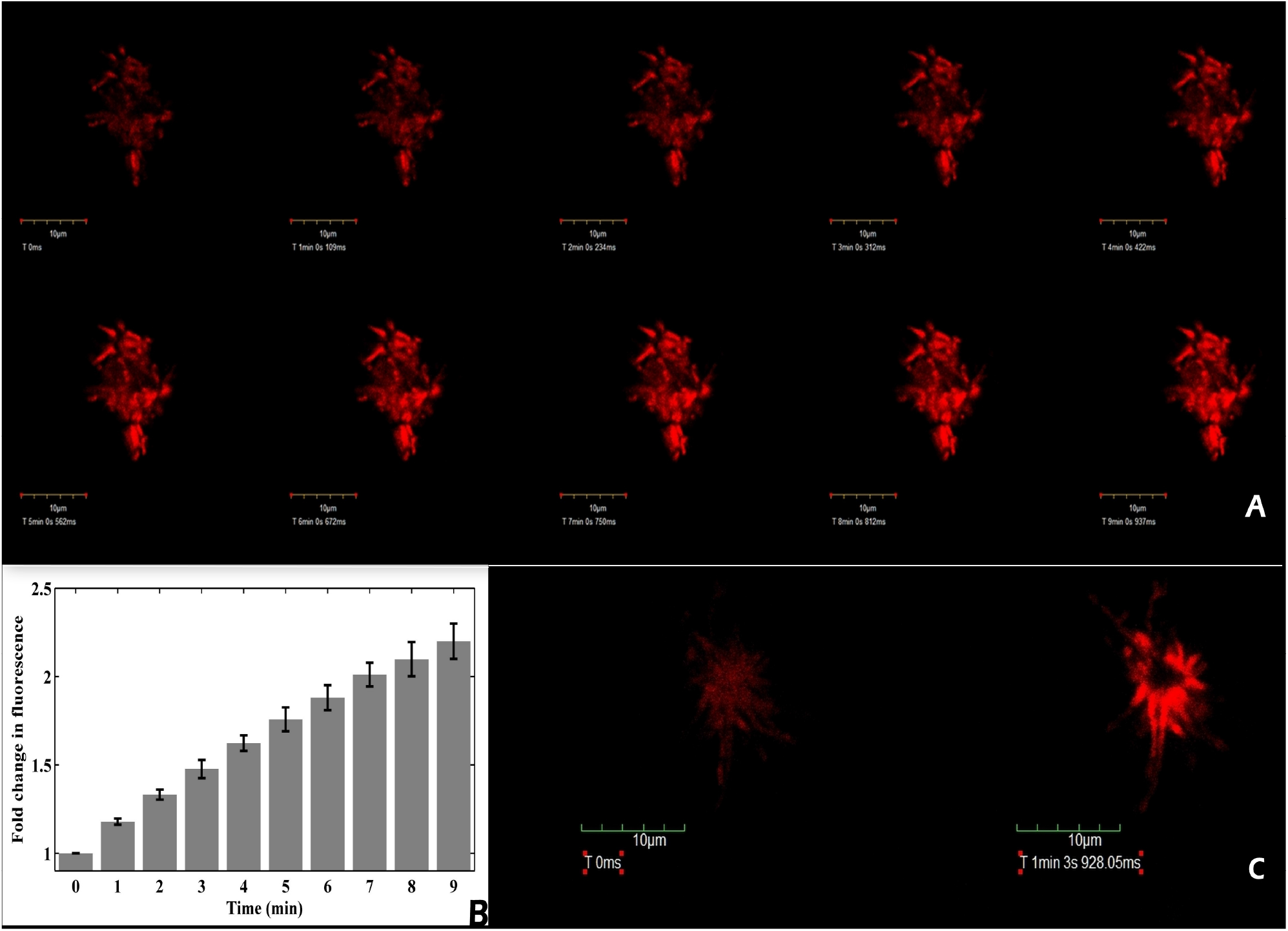
Panel A:Shows gradual increase in fluorescence emission of bacterial cell cluster. Cells were exposed to 405 nm laser (25%intensity) up to 9’, at 1’ interval starting from 0’. The lower panel B shows the relative increase in image intensity (with respect to 0 min, the error being evaluated using four indepndent ROIs). Panel C shows image of a cell cluster exposed for 60 s to 405 nm laser (100%intensity); The initial image (left) after 60 seconds of laser exposure assumes the image form shown at the right (of the panel C). Whereas, the central zone of the right image is bleached due to intense laser exposure a more illuminated zone appears at some distance.

### Spectroscopic studies

Absorbance spectra of the cell free extract (see figure() 2)) show a soret band at 395 nm and Q band at 500 nm, 535 nm and 560 nm. The pattern of absorbance spectra indicates the presence of H and J band respectively near 400 nm and 535 nm (approximate ratio 7:1). The absorbance value of the sample shows a sustained increase with time of UV exposure.

**Figure 2:**
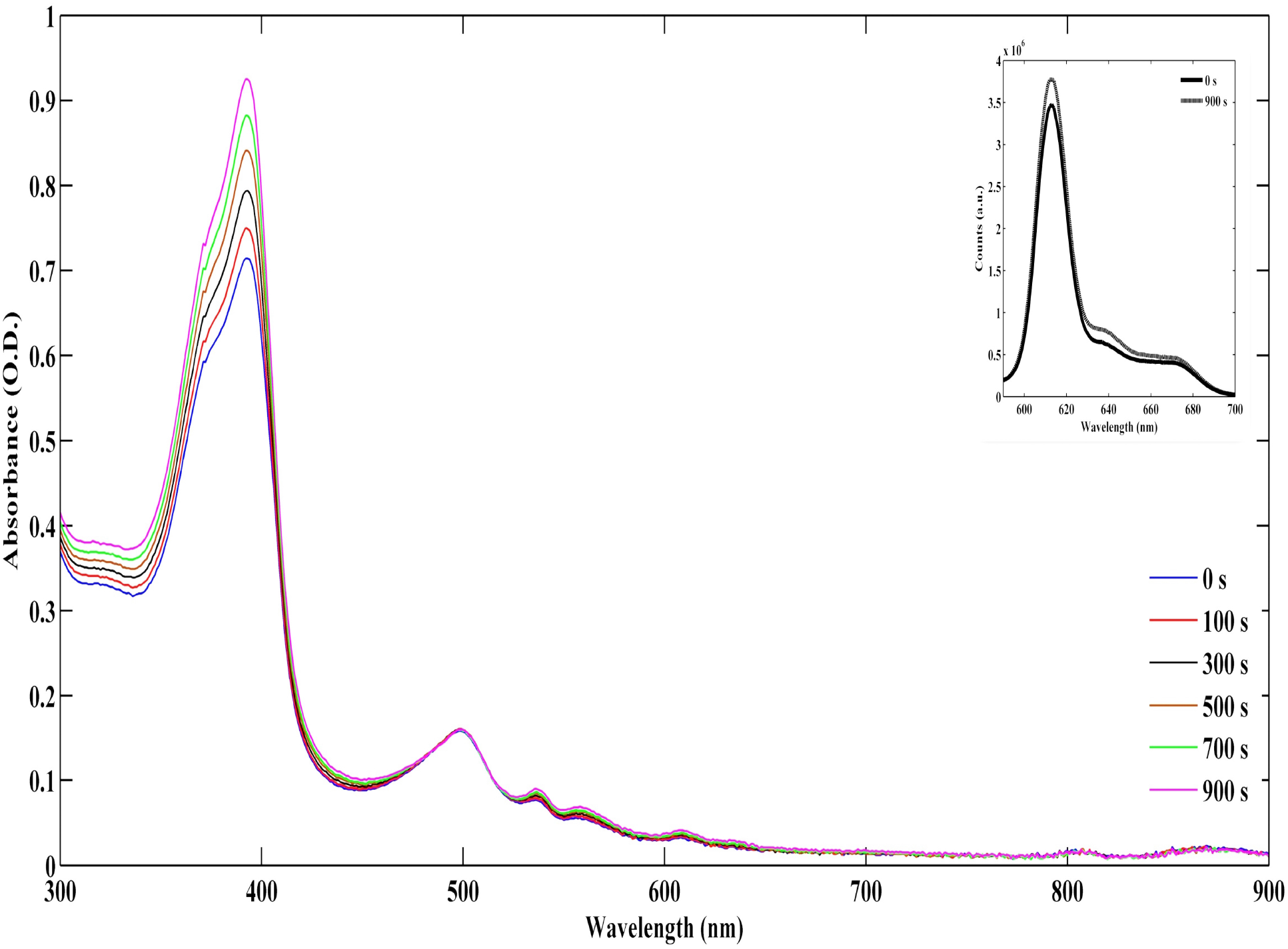
The absorbance spectra of the cell free extract shows gradual increase in absorbance at 395 nm upon prolonged excitation (concentration remaining identical). The inset compares initial (solid line) and final (dashed line) fluorescence spectra indicating amplification in fluorescence upon such excitation.

The system was found to fluoresce near 613 nm when excited at λ_*max*_ = 395 nm. The bacterial cells and the cell free extract showed similar fluorescence properties (see figure (S2a)).

The time based fluorescence upon excitation at 395nm is shown in figure(S2a). The time scan reveals an increase in fluorescence emission intensity with time for both whole cells and the cell free extract. The rate of amplification in the two cases are however different. When acetone was selected as solvent medium two additional peaks appeared at 594nm and 635nm. Notably, the excitation is also red shifted to 411nm and 403 nm for the respective emissions. The split of original spectral pattern (see figure(S2a)) into two peaks reveals the fact that the integrity of the membrane bound complex is sensitive to solvent perturbation. Notably, only one peak (635nm) among the two, is amplified with time when there is a prolonged excitation (see dotted profile in the figure (S3)). There is some variations of exact peak position depending on the bacterial growth conditions, the robust result remaining similar.

### Concentration dependent bistability

The cell free extract of bacterial culture shows transition in emitted fluorescence intensity over a threshold concentration. Figure (3) reveals concentration dependent enhanced emission during a fluorescence time scan. One can easily follow fluorescence emission intensity either rises with time or decreases with time depending on concentration of the fluorophore. For further insight of such concentration dependent enhanced emission, we studied the initial (first 20 s) fluorescence emission rate.

**Figure 3:**
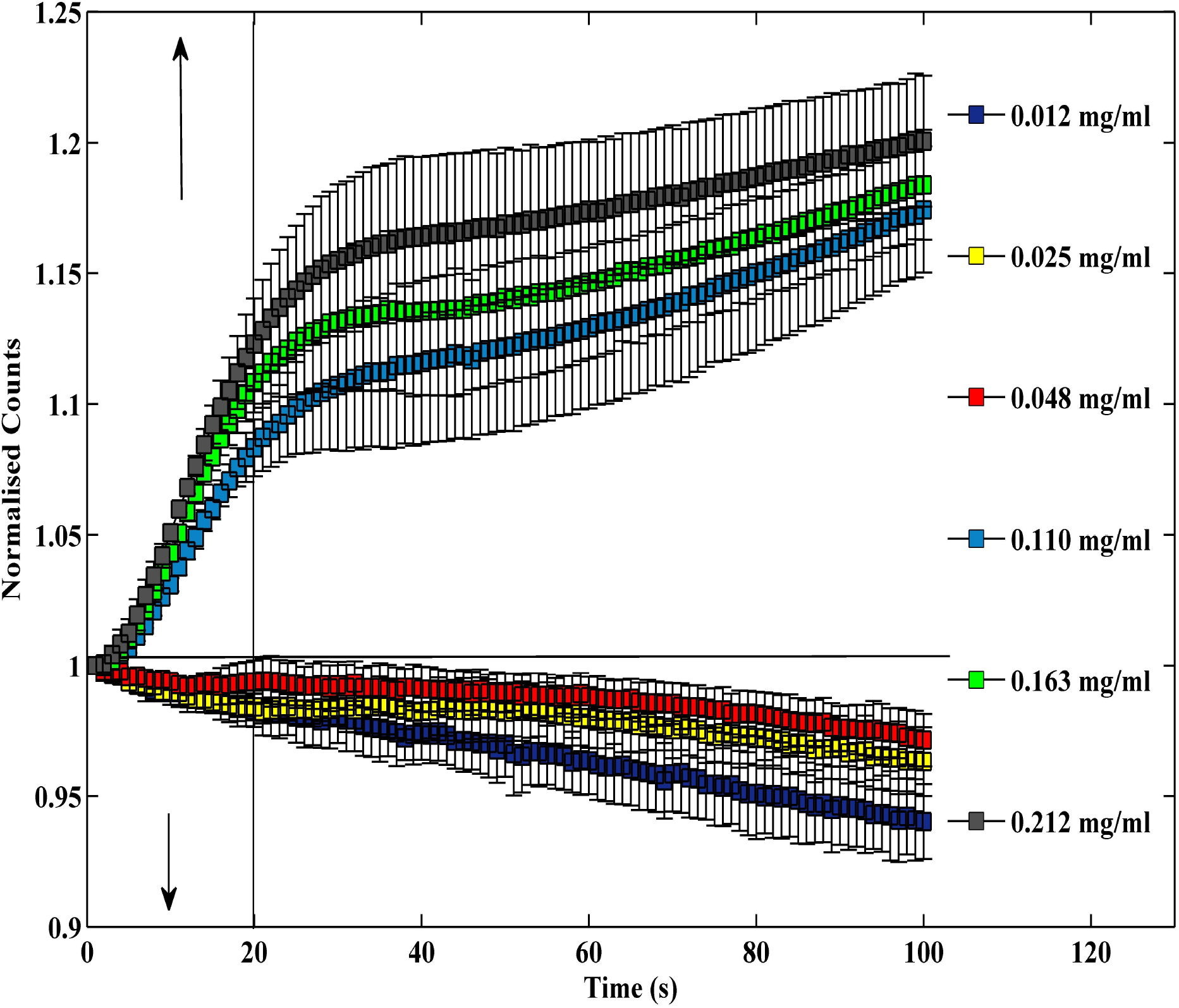
Figure shows how changing the fluorophore concentration switches the staeady states from bleaching to amplifying states of fluorescent emission. Time based fluorescence emission normalized with initial fluorescence intensity along Y axis is plotted with concentration along X axis, error level being represented by the vertical lines. The bistability occurs between the critical concentration below 0.048 and above 0.11 *mg · ml*^−1^.

Rate of fluorescence emission (change in fluorescence emission intensity at maximum emission wavelength (counts) with per unit time) was plotted against function of concentration of the fluorophore (see figure(4)) where the concentration of the fluorophore is expressed by absorbance at 395 nm (the excitation maxima of the fluorophore). Figure(4) reveals that not only fluorescence intensity but also the amplification has a systematic dependence on the concentration; rate of emission changing from positive to negative with dilution of the fluorophore concentration.

**Figure 4:**
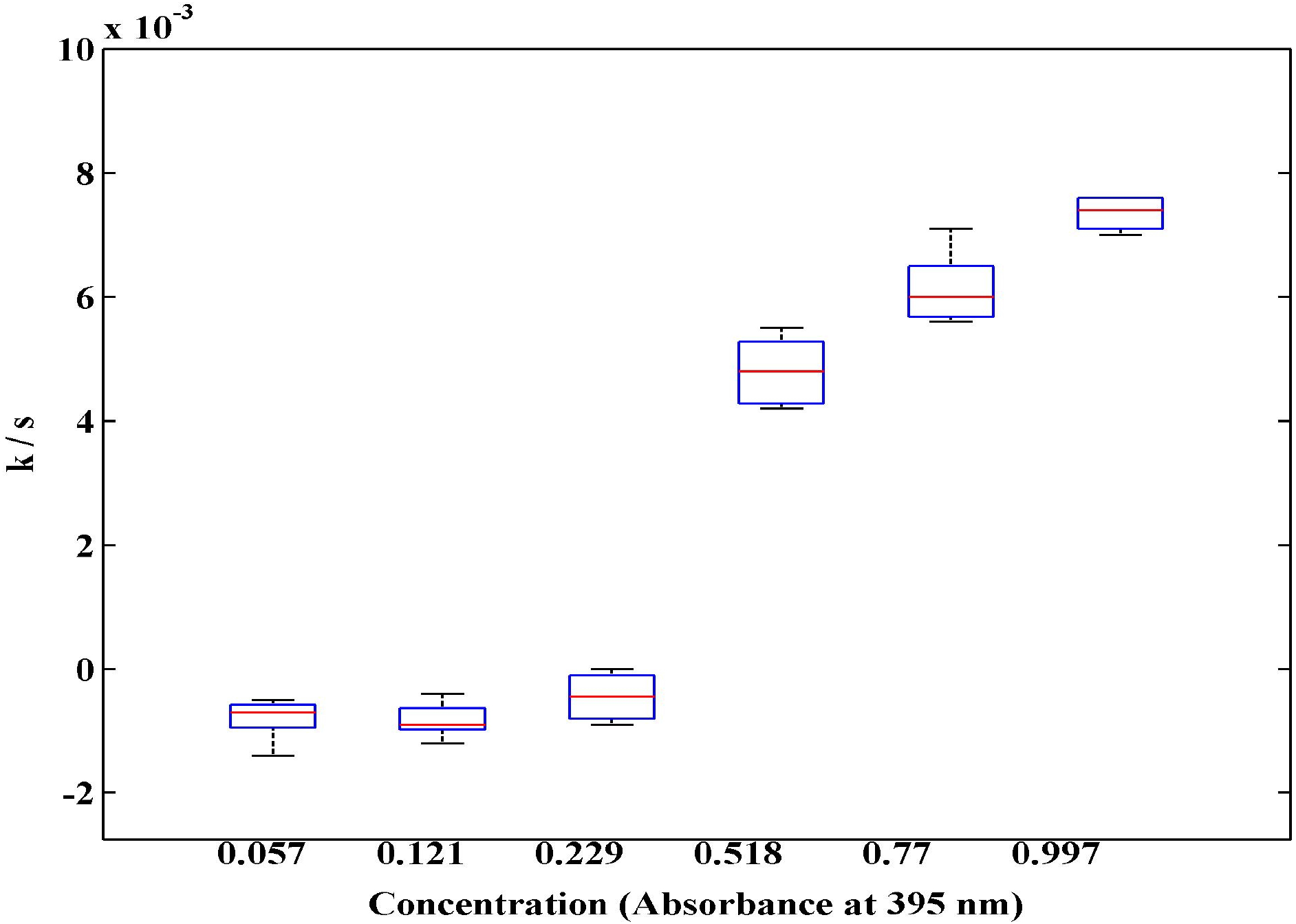
Fluorescence amplification rate expressed by k(*s*^−1^) along Y axis is plotted against concentration(X 0.1 *mg · ml*^−1^) along X axis.

### Temperature dependence and static magnetic field effect

The cell free extract of higher concentration (O.D. 0.1 at 395 nm) shows a systematic decrease in fluorescence counts at 278K in contrast to 298K (see upper panel of figure (S2b)). But when the sample concentration is further diluted to ten times (O.D. 0.01 at 395 nm) shows no difference with temperature change. In lower panel of figure(S2b) the time based fluorescence of the above samples is shown. figure(5) illustrates the effect of Static Magnetic Field (SMF) on fluorescence property of the system. The fluorescence amplification process was found best at 298K. But at low temperature (below 280K), the amplification process is attenuated (see upper panel of figure(5)). The comparison of upper and lower panels of figure(5) clearly indicates that at lower temperature, the amplification reappears in presence of SMF.

**Figure 5:**
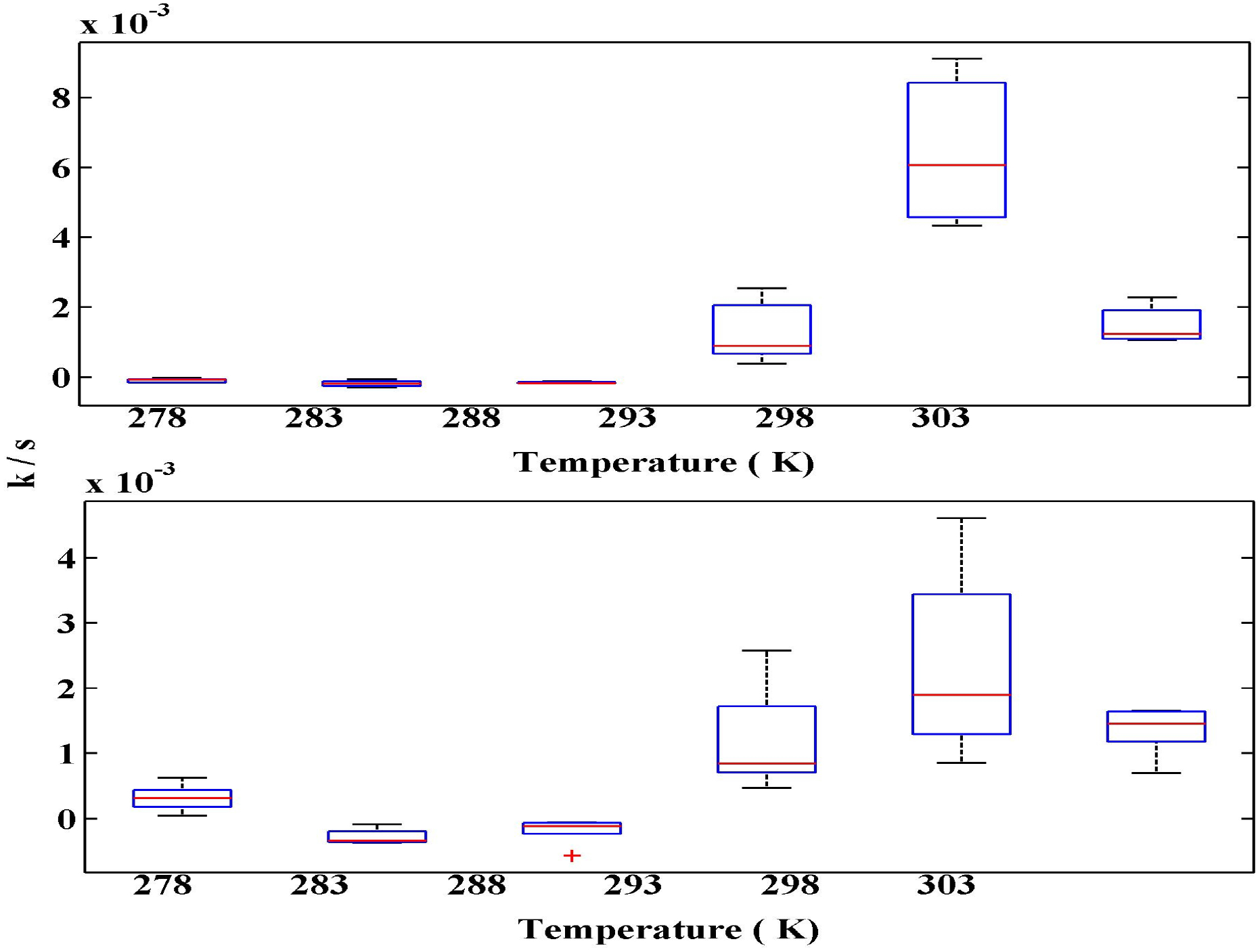
Fluorescence amplification rate at 613 nm (k s^−1^) for excitation at 395 nm as function of T(in °*K*) in absence (upper panel) and presence (lower panel) of magnetic field.

### Fluorescence Life time

Life time studies revealed a few characteristics of the amplitude gain phenomenon. Dilution of sample with solvent shows that the life time value decreases with increasing order of dilution, higher dilutions have low life time values (see Table 1).

**Table 1:** Life time as a function of fluorophore concentration

| Conc.mg · ml<sup>-1</sup> | Life time (ns) |
| --- | --- |
| 0.039 | 1.78, 15.7 |
| 0.113 | 1.90, 15.0 |
| 0.590 | 2.59, 15.6 |
| 1.206 | 2.74, 15.6 |
| 2.984 | 2.99, 15.6 |

The evaluation of life time values for the excreted fluorophore revealed order of 2 nanoseconds and 15 nanoseconds which indicate formation of two species in excited states. Where the shorter one is prone change life time upon perturbation of surrounding environment which could be due to alteration of structural changes on the other hand the longer one is more or less static. So, the notable point is that during fluorescence emission there should be a simultaneous competition between two upon excitation for long time as one species is found to show varying lifetime and another is not.

### Dynamic Light Scattering (DLS) measurement

At low temperature (278K) the sample shows abundance of small aggregates which is again shifted to lager population size when exposed to a static magnetic field (see Fig. 6). DLS measurement of the cell free extract shows temperature induced aggregation of the sample. In figure (7) when mean intensity was plotted against the hydrodynamic size it was found that with increasing temperature there is a tendency of larger population size with heterogeneous aggregates. from Table 3 it is clearly observed that The Z average value of the sample is increasing with temperature. At 278K the sample shows a increased Z average value in presence of SMF though the poly Dispersity Index is also increased. At 298K, 303K and 313K a very small population arises with respectively small hydrodynamic size which is absent in 278K, 288K and 313K (see Fig. 7 and Fig. 6).

**Figure 6:**
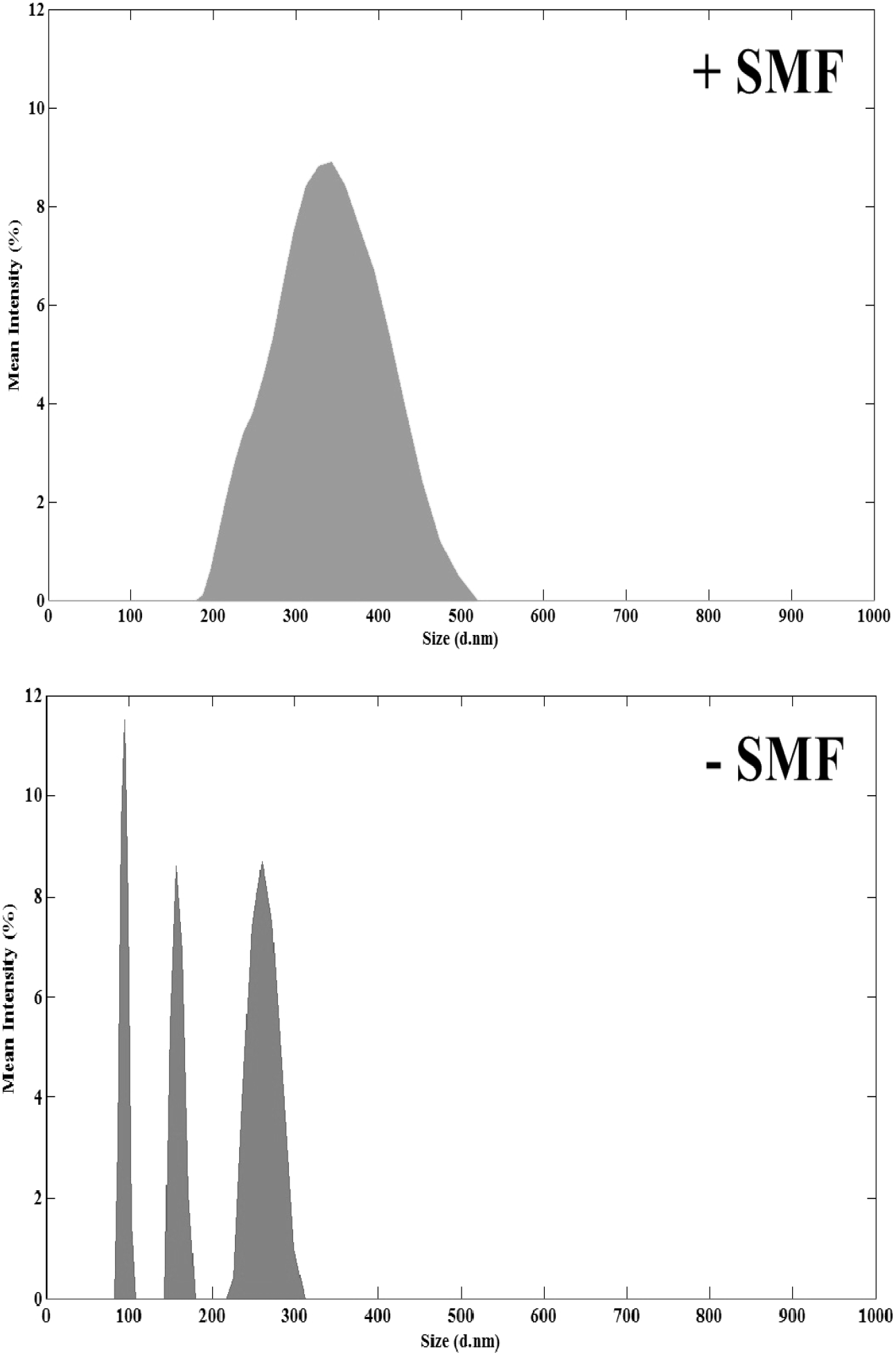
Dynamical light scattering measurement showing diversity of aggregate morphology in presence and absence of static magnetic field at 278K.

**Table 2:** OR gated T and B dependent bifurcations

| T>278K (O1) | B (SMF 0.25 T) (O2) | Amplification (A) |
| --- | --- | --- |
| 0 | 0 | 0 |
| 0 | 1 | 1 |
| 1 | 0 | 1 |
| 1 | 1 | 1 |

**Table 3:** T dependence of Polydispersity Index (PDI)

| Temperature | SMF (0.0285 T) | Z Average (d.nm) | Average PDI |
| --- | --- | --- | --- |
| 278K | — | 356.8 ± 46.47 | ≤ 1 |
| 278K | + | 828 ± 76.38 | < 0.5 |
| 288K | — | 849.7 ± 41.43 | ≤ 1 |
| 298K | — | 758.1 ± 36.38 | ≤ 0.5 |
| 303K | — | 755.1 ± 161.5 | < 1 |
| 313K | — | 997.1 ± 36.20 | ≤ 1 |

**Figure 7:**
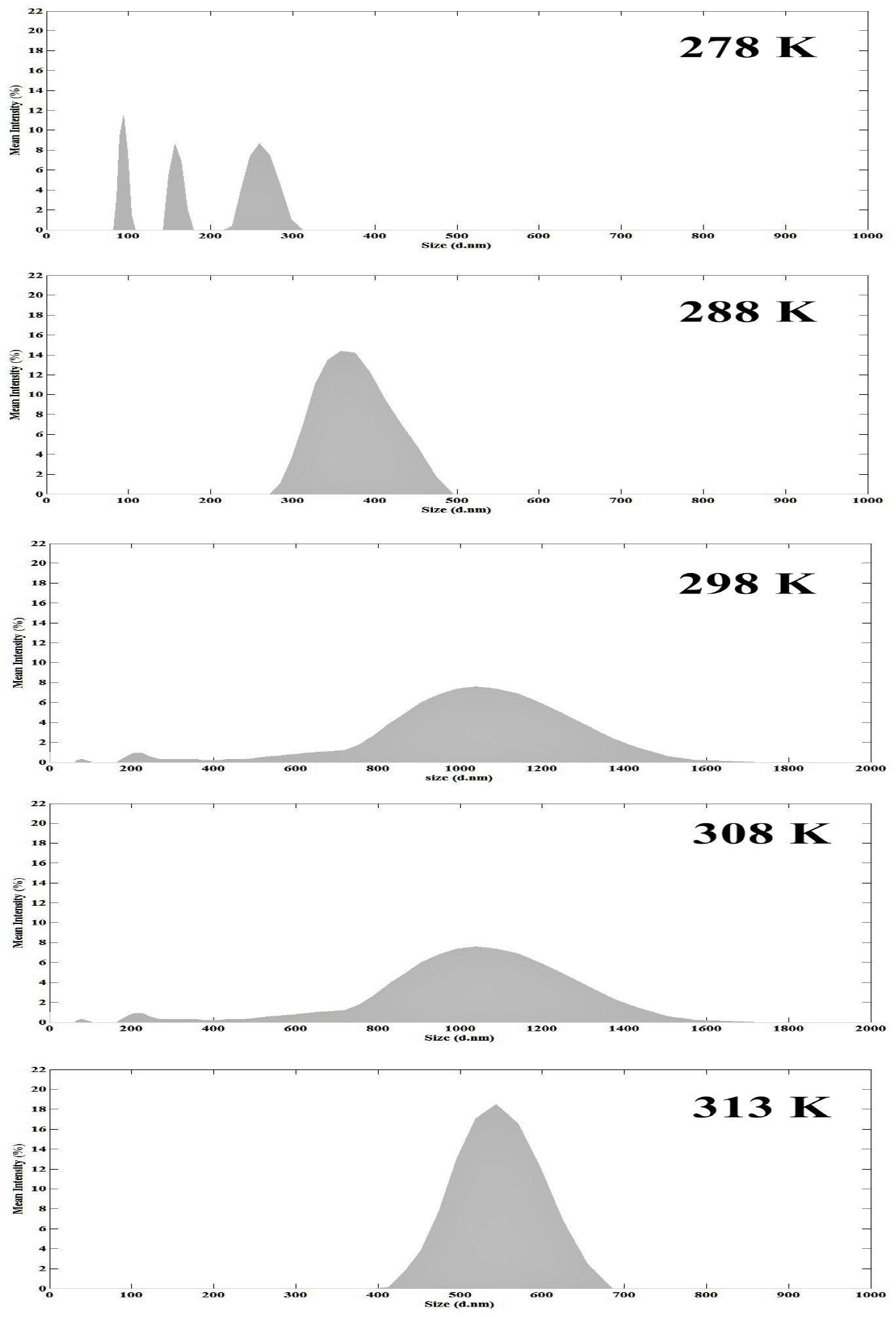
Temperature dependence of aggregate formation, higher temperature showing aggregates with higher hydrodynamic radius.

## Discussion

The bistability in dynamical system is not a concurrent existence of two steady states. It rather implies possibility of transition from one steady state to another, in response to variations of a critical parameters(see the gate like situation summarized in Table (2)).The co-operative change of the kinetic constant with respect to variations in concentration (figure (4)) & temperature (figure(5)) suggests onsets of criticality in concentration or tempeartaure dependent transitions. The striking feature of the reported bistability in *Rhodobacter capsulatus* is that more than one critical parameters lead to bifurcation behavior.

In simpler terms, there seems to be numerous parameters whose change can induce critical bistable transitions of the optical emission status from fluorescence quenching to amplification states or *vice versa*. For example, increase in population density triggers such transition. If temperature is lowered a reverse transition occurs resulting in quenching. A reverting back to amplifying state occurs if we introduce static magnetic field.

What seems intriguing is the confocal microscopic data figure (1). The panel A and the intensity variation shown in panel B of this figure matches with the spectroscopic analysis (see upper panel of figure(3)). In both cases gradual enhancement of fluorescence at moderate fluorophore concentration are suggested. The panel C in figure(1) however needs additional analysis. It follows that excitation intensity can serve another bifurcation parameter, higher intensity (in the central part) causing quenching, whereas moderate intensity (in the peripheral region) showing amplification. Dead microbial cells in which biofilm formation is complete (data not shown) does not exhibit amplification. The porphyrin excretion and their photon induced aggregation thus seems to be an attribute of the live microbes and the chance of their use in cell cell communication seems imminent. The aggregation may thus constitute a component of photon sensing machinery for this class of photosynthetic bacteria.

The optical sensing by the functional microbial population is actually a rational combination of photon excitation and protoporphyrin aggregate formation. The aggregates can enhance the fluorescent yield.

If *Rhodobacter capsulatus* cells are grown under anaerobic or micro-aerophilic condition in presence of continuous light they do accumulate tetrapyroles (Protoporphyrin, Mg-Protoporphyrin derivatives) (27). We can postulate aggregation of such natural porphyrin derivatives are synthesized by photosynthetic bacteria. Porphyrins contain tetrapyrroles which could easily interact non-covalently because of the presence of *π*bonds, (28, 29). While self assembly is known to be favored at low temperature, (30)for porphyrins, they undergo aggregate formation with increasing temperature (31). The aggregate normally has a higher quantum yield for fluorescence and the observation that amplification occurs at moderate and not at low temperature is supportive of the fact that what we are observing in course of amplification is aggregation of porphyrin. Why low temperature shows magnetic sensitivity can be explained by extrapolating this arguments and additionally postulating that a different class of aggregation occurs at lower temperature. That low temperature facilitates a differential aggregate formation is supported by figure (7). The magnetic susceptibility of the aggregates formed at low temperature (see figure (7)) may be a reflection of the fact that over all spin states may have critical dependence on aggregation.

Porphyrins with negatively charged peripheral groups easily aggregate in aqueous solution (32, 33). At room temperature, porphyrins could coexist in monomeric and aggregated forms. The monomers are however reactive to oxygen and this could be one of the reasons for porphyrin accumulation by cells as a future defense to survive aerobic condition. According to the present literatures porphyrins may form J and H type of aggregates (34, 35) and they could remain in a mixture of the two (36) in room temperature. One of the aggregates are indeed sensitive to magnetic field as indicated earlier. If we conjecture that at low temperature there is a shift of equilibrium towards the J-aggregates the results completely matches with our observation. The cooperative nature of the concentration dependence implies higher concentration of monomers actually facilitates the aggregate formation by distributing the incident light. With increasing concentration the equilibrium is thus shifted to higher aggregate formation and the rate of fluorescence enhancement increases. In higher concentration the fact is that the molecular crowding favours photon induced aggregation which in turn could explain the co-operative concentration dependent rise in fluorescence (see the sigmoidal nature of figure (4)). Eventually magnetic sensitivity of certain classes of stacked pi ring assemblies was reported by our group earlier (37) (also see Fig. 6). Investigation of the ecological aspect of such phenomenon could consider the fact that the heterogeneous large aggregation of the porphyrin derivatives (39) in aqueous medium should provide matrix for the cells to form biofilm structure.

## Conclusion

Photon capture and its subsequent bio-energetic management is in the bottom of the ecological pyramid. We demonstrate multiplication of fluorescence emission in the red region for excitation at near UV region by a photosynthetic bacterium *R.capsulatus*. A thermal switch puts off the multiplicative emission at lower temperature. Such low temperature driven repression of the emission gain is reversed in presence of a moderate dose of static magnetic field. The work suggests that one can employ the microbe as Maxwell’s demon to minimize ecological damage caused by UV, as it absorbs UV and emits photochemically inert red light. The porphyrin aggregate (see figure S5) formation may explain the logical and bifurcating fashion by which the demon may actually work. We must mention the ecological significance of the result. It is intriguing to note that the principle of population inversion on which the lasers are made is exploited by one of the ancient life forms. The amplification of fluorescence can serve as a photonic agent of quorum sensing though the mechanism behind suggests natural defense to survive the effect of oxygen in growth medium (40) or to compete with other species when exposed to a heterogeneous population(41).

The practical use of this concentration, temperature and magnetic field sensitive system is evident and it may be possible to construct synthetic biological circuits for addressing issues related in the field of aggregation induced enhanced emission systems(42) by appropriate extrapolation of the above.

## Acknowledgment

We thank Department of Biotechnology, Govt. of India for supporting the research (grant number BT/PR3957/NNT/28/659/2013). We thank Dr. Shibsekhar Roy, Dr.Swagata Banerjee and Dr. Jaydeep Bhattacharya for helpful suggestions and Ms. Boni Haldar, Ms. Puja Biswas (DBT IPLS) for technical assistance in confocal and AFM experiments. We also thank Prof. M.Bhattacharya for providing the life time measurement facility.

